# AlphaFold2-aware protein-DNA binding site prediction using graph transformer

**DOI:** 10.1101/2021.08.25.457661

**Authors:** Qianmu Yuan, Sheng Chen, Jiahua Rao, Shuangjia Zheng, Huiying Zhao, Yuedong Yang

## Abstract

Protein-DNA interactions play crucial roles in the biological systems, and identifying protein-DNA binding sites is the first step for mechanistic understanding of various biological activities (such as transcription and repair) and designing novel drugs. How to accurately identify DNA-binding residues from only protein sequence remains a challenging task. Currently, most existing sequence-based methods only consider contextual features of the sequential neighbors, which are limited to capture spatial information. Based on the recent breakthrough in protein structure prediction by AlphaFold2, we propose an accurate predictor, GraphSite, for identifying DNA-binding residues based on the structural models predicted by AlphaFold2. Here, we convert the binding site prediction problem into a graph node classification task and employ a transformer-based variant model to take the protein structural information into account. By leveraging predicted protein structures and graph transformer, GraphSite substantially improves over the latest sequence-based and structure-based methods. The algorithm is further confirmed on the independent test set of 181 proteins, where GraphSite surpasses the state-of-the-art structure-based method by 16.4% in AUPR and 11.2% in MCC, respectively. We provide the datasets, the predicted structures, and the source codes along with the pre-trained models of GraphSite at https://github.com/biomed-AI/GraphSite. The GraphSite web server is freely available at https://biomed.nscc-gz.cn/apps/GraphSite.

## 1. Introduction

Protein-DNA interactions play crucial roles in many biological processes such as transcription, repair, and signal transduction [1, 2]. While protein-DNA binding affinity prediction [3, 4], protein-interacting site prediction on DNA (e.g. promoter prediction) [5], and protein-DNA docking [6] have been widely studied, accurately identifying amino acids involved in protein-DNA interactions solely based on proteins is also an important topic in bioinformatics, which helps to improve molecular docking [7, 8], understand disease mechanism [9, 10], predict protein function [11, 12], and identify potential drug target for novel drug design [13, 14]. However, conventional experimental methods for DNA-binding site detection such as X-ray crystallography [15] and fast ChIP [16] are costly and time-consuming. Although protein-DNA binding patterns are complicated and many proteins may specifically recognize local DNA structures such as hairpins, cruciforms and G-quadruplexes [17, 18], DNA-binding residues are often conserved [19]. Therefore, it is necessary and feasible to develop complementary computational methods capable of making reliable and accurate DNA-binding site prediction.

Computational methods for DNA-binding site prediction can be classified into two classes, sequence-based and structure-based methods, according to their used information. Sequence-based methods such as DNAPred [20], DNAgenie [21], and NCBRPred [22] learn local patterns of DNA-binding characteristics through sequence-derived features. For example, NCBRPred adopts evolutionary conservative information and predicted secondary structure and solvent accessibility extracted from protein sequence and employs bidirectional Gated Recurrent Units (BiGRUs) to learn local patterns from sequence contexts using sliding-window strategy. While these sequence-based approaches require protein sequences only, the lacks of using tertiary structures cause their limited predictive accuracies.

By comparison, structure-based approaches inferring binding sites from known structures are often more accurate, which can be generally categorized into template-based methods, machine learning based methods, and hybrid methods. Template-based methods identify DNA-binding sites using the sequence and structure information of templates, which are selected by alignment or comparison algorithms [23, 24]. Nevertheless, for the proteins that have no high-quality template, the performance of these methods will be seriously restricted. With features derived from protein structures, recent structure-based machine learning methods represent protein structures as voxels in three-dimensional Euclidean space or nodes in connected graphs. For example, DeepSite [25] maps protein atoms into 3D voxels and employs 3D convolutional neural networks (3DCNN) to extract features from neighborhood of the target residue. Alternately, GraphBind [26] encodes protein structures as graphs and adopts graph neural networks (GNN) to learn the local tertiary patterns for binding residue prediction. Hybrid methods integrate template-based methods and machine learning based methods simultaneously, such as DNABind [27], COACH-D [28] and NucBind [7]. Albeit powerful, the structure-based methods are not applicable to most proteins that don’t have known tertiary structures due to the difficulties to determine protein structures experimentally [29].

With the development of deep learning techniques, protein structure prediction is experiencing a breakthrough. The representative method, AlphaFold2 [30], has incorporated physical and biological knowledge about protein structure, information of multi-sequence alignment, and the sophisticated design of the deep learning algorithm. The method was shown able to predict protein structure with atomic accuracy even when no similar structure template is known, and demonstrated accuracy competitive with experiment in a majority of cases in the challenging 14th Critical Assessment of protein Structure Prediction (CASP14). Such breakthrough will undoubtedly benefit downstream protein function studies, including binding site prediction.

Effective learning of protein structure remains a challenging task, even though 1DCNN [31], 2DCNN [32], 3DCNN [33], GNN and its variants [34, 35] have been widely adopted. On the other hand, transformer [36] is well acknowledged as the most powerful neural network in modelling sequential data, such as natural language[37], drug SMILES [38] and protein sequence [39]. In the last few years, transformer variants have also been shown great performance in graph representation learning [40–42]. Therefore, it is promising to advance the protein binding site prediction by constructing accurate structure model from sequence and effectively learning the structural information through the recent graph transformer technique.

In this study, we have developed a novel method GraphSite, which applies graph transformer network and predicted protein structures from AlphaFold2 for sequence-based prediction of DNA-binding residues. Specifically, we integrate multi-sequence alignment and structural information to construct residual features and calculate pairwise amino acid distances to mask out the spatially remote amino acids when calculating attention scores in the transformer. With the spatial information and structure-aware transformer, GraphSite was found to outperform other sequence-based and structure-based methods through various performance evaluations. To the best of our knowledge, this is the first work that utilizes AlphaFold2-predicted structures and graph transformer for protein-DNA binding site prediction, which can be easily extended to sequence-based prediction of other functional sites.

## 2. Materials and Methods

### 2.1 Datasets

We adopted two publicly available benchmark datasets from the previous study [26] to train and test our method: Train_573 and Test_129, which are named by the numbers of proteins in the datasets. These two datasets were collected from the BioLiP database [43], which pre-computes binding sites according to experimentally determined complex structures from Protein Data Bank (PDB) [44]. Concretely, Train_573 contains proteins released before 6 January 2016 while Test_129 from 6 January 2016 to 5 December 2018. In these datasets, a DNA-binding residue was defined if the smallest atomic distance between the target residue and the DNA molecule is less than 0.5 Å plus the sum of the Van der Waal’s radius of the two nearest atoms. To deal with the data imbalance problem, the authors [26] applied data augmentation on Train_573, which transferred binding annotations from protein chains with similar sequences (sequence identity > 0.8) and structures (TM scores > 0.5) to increase the number of binding residues. This was conducted for the following reasons: 1) similar proteins, although could derived from different organisms, may have the same biological function. 2) different resolutions or co-factors may lead to minor differences in the structures for the same protein. Finally, CD-HIT [45] was used to ensure no redundant protein with more than 30% sequence identity within the training set and between the training and test set. To further demonstrate the generalization of our method, we built another independent test set (Test_181) based on newly released DNA-binding proteins in BioLiP (6 December 2018 to 19 August 2021). We have removed redundant proteins sharing sequence identity > 30% over 30% overlap with any sequence in the above two datasets and within Test_181 using CD-HIT. Details of the number of binding and non-binding residues of these datasets are given in **Table 1**.

**Table 1.** Statistics of the three benchmark datasets used in this study. The columns give, in order, the dataset name, the number of binding, and the number of non-binding residues in each dataset, and the percentage of the binding residues out of total.

| Dataset | Binding residues | Non-binding residues | % of binding residues |
| --- | --- | --- | --- |
| Train_573 | 14479 | 145404 | 9.06 |
| Test_129 | 2240 | 35275 | 5.97 |
| Test_181 | 3208 | 72050 | 4.26 |

### 2.2 Protein representation

In our framework, the DNA-binding site prediction task is treated as a graph node classification problem, where a protein consisting of *n* amino acid residues is represented by a node feature matrix ***X*** and a distance matrix ***D***.

#### 2.2. Predicted protein structures and distance maps

Following the tutorial at https://github.com/deepmind/alphafold, we downloaded the model parameters and genetic databases including UniRef90 [46], MGnify [47], BFD [48], Uniclust30 [49], PDB70 [50] and PDB [44] to implement AlphaFold2 in the Tianhe-2 supercomputer. We set the “--max_template_date” parameter to 2020-05-14 just as the AlphaFold2 model in CASP14 to predict the protein structures in Train_573 and Test_129, from which the relaxed models with the highest confidences (measured by the predicted Local Distance Difference Test (LDDT) scores) were chosen. However, to further avoid the possibility that AlphaFold2 might use the native known structures in PDB as templates, we set “--max_template_date” to 2018-12-5 when predicting proteins in the independent Test_181. This made Test_181 more challenging and closer to the situation where only protein sequences instead of similar templates are available. According to the predicted protein structural models from AlphaFold2, we acquired the coordinate of the Cα atom of each amino acid residue, and then calculated the Euclidean distances between all residue pairs, which formed a distance map 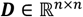.

#### 2.2.2 Node features

We employed two groups of amino acid features to train our model: multi-sequence alignment (MSA) information and structural properties, which were concatenated and formed the final node feature matrix 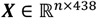 with *n* denoting the length of a protein sequence.

##### MSA information

Co-evolving amino acids may structurally contact, and evolutionarily conserved residues may contain motifs related to important protein properties such as DNA binding propensity. Here, we employed the “single representation” feature output by AlphaFold2 along with the predicted structures. This single representation is derived by a linear projection of the first row of the MSA representation, which is a highly processed MSA feature matrix through 48 Evoformer blocks in AlphaFold2. In addition, we also explored the widely used position-specific scoring matrix (PSSM) and hidden Markov models (HMM) profile. Concretely, PSSM was generated by running PSI-BLAST [51] to search the query sequence against UniRef90 database with three iterations and an E-value of 0.001. The HMM profile was produced by running HHblits [52] to align the query sequence against UniClust30 database with default parameters. Each amino acid was encoded into a 384-dimensional vector in single representation and 20-dimensional vector in PSSM or HMM, and the values were normalized to scores between 0 to 1 using Equation (1), where *v* is the original feature value, and *Min* and *Max* are the smallest and biggest values of this feature type observed in the training set.

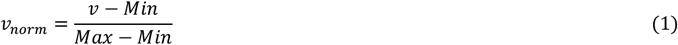

##### Structural properties

Three types of structural properties were extracted by the program DSSP [53] using predicted structures: 1) 9-dimensional one-hot secondary structure profile where the first 8 dimensions represent 8 secondary structure states, and the last dimension represents unknown secondary structure. 2) Peptide backbone torsion angles PHI and PSI, which were converted to a 4-dimensional feature vector using sine and cosine transformations. 3) Solvent accessible surface area (ASA), which was normalized to relative solvent accessibility (RSA) by the maximal possible ASA of the corresponding amino acid type. This 14-dimensional structural feature group is named DSSP in this article.

### 2.3 The architecture of GraphSite

**Figure 1** shows the overall architecture of the proposed framework GraphSite, where the protein sequence is input to AlphaFold2 to produce the single representation and the predicted protein structure, from which the distance map and DSSP are extracted. Finally, the single representation, DSSP and sequence-derived features PSSM and HMM are concatenated to form the node feature vector, which is then input to the graph transformer masked by the distance map to learn the DNA-binding site patterns.

**Figure 1.**
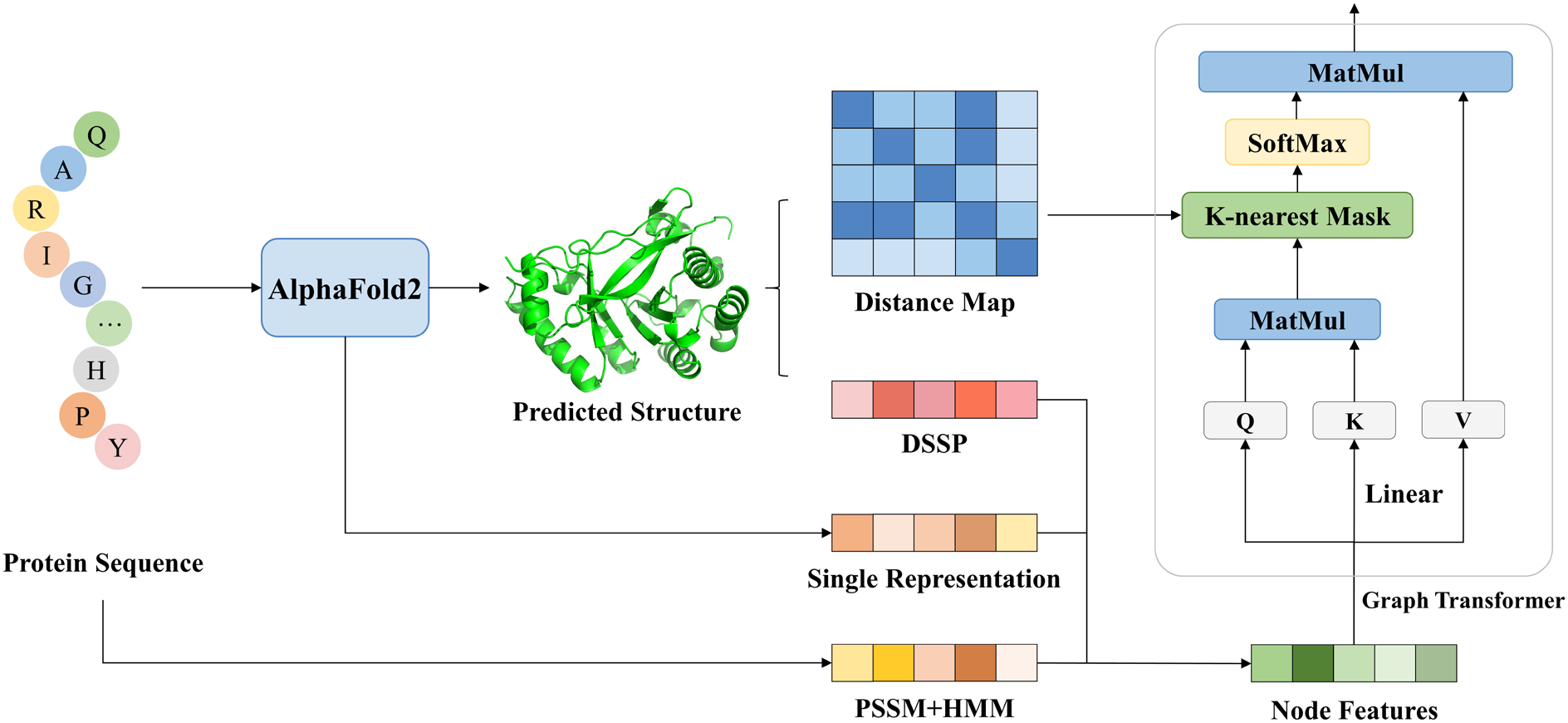
The overall architecture of GraphSite. The protein sequence is input to AlphaFold2 to produce the single representation and the predicted protein structure, from which the distance map and DSSP are extracted. Then, the single representation, DSSP and sequence-derived features PSSM and HMM are concatenated to form the node feature vector, which is then input to the graph transformer model with *k*-nearest mask by the distance map to learn the DNA-binding site patterns.

#### 2.3.1 Graph transformer

The traditional transformer encoder layer consists of a multi-head self-attention module and a position-wise feed-forward network. Let 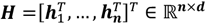 denote the input of the self-attention module where *d* is the hidden dimension and 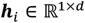 is the hidden representation of the *i*^th^ amino acid. Note that the initial ***H***^(0)^ does not have to be the feature matrix ***X***, which means that a fully connected layer can be applied on ***X*** to obtain the initial hidden representation ***H***^(0)^. The input ***H***^(l)^ is projected by three matrices 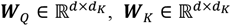 and 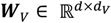 to the corresponding queries, keys and values representation ***Q, K, V***:

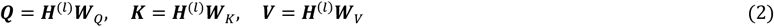

The self-attention is then calculated as:

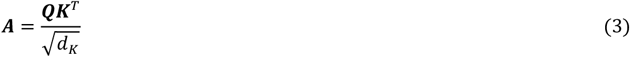

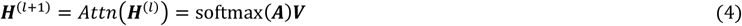

where ***A*** is a matrix capturing the similarities between queries and keys. In order to take the protein structure information into account for focusing on spatially adjacent residues, we adopt *k*-nearest mask according to the distance matrix ***D*** to mask out the spatially remote amino acids, which means that for node *i*, only spatially adjacent nodes *j* ∈ Neighbor(*i, k*) are used to calculate the attention scores in the transformer. In order to jointly attend to information from different representation subspaces at different positions, we use multi-head attention to linearly project the queries, keys and values *h* times, perform the attention function in parallel and finally concatenate them together. In this study, *d_K_* = *d_V_* = *d* / *h*.

#### 2.3.2 Multilayer Perceptron

The output of the last graph transformer layer is input to the multilayer perceptron (MLP) to predict the DNA-binding probabilities of all *n* amino acid residues:

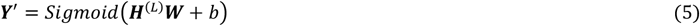

where 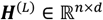 is the output of the *L*^th^ graph transformer layer; 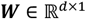 is the weight matrix; 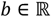 is the bias term, and 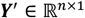 is the predictions of *n* amino acid residues. The sigmoid function normalizes the output of the network into binding probabilities ranging from 0 to 1.

### 2.4 Implementation details

We performed 5-fold cross-validation on the training data, where the data were split into five folds randomly. Each time, a model was trained on four folds and evaluated on the remaining one fold. This process was repeated five times and the performances on the five folds were averaged as the overall validation performance, which was used to choose the best feature combination and optimize all hyperparameters through grid search (**Supplementary Table S1**). In the testing phase, all five trained models in the cross-validation were used to make predictions, which were averaged as the final predictions of our method.

Specifically, we utilized a 2-layer graph transformer module with 64 hidden units and the following set of hyperparameters: *h* = 4, *k* = 30 and batch size of 16. We employed the Adam optimizer [54] with *β_1_* = 0.9, *β_2_* = 0.99, *ε* = 10^−5^, weight decay of 10^−5^ and learning rate of 3 × 10^−4^ for model optimization on the binary cross entropy loss. The dropout rate was set to 0.2 to avoid overfitting. We implemented the proposed model with Pytorch 1.7.1 [55]. Within each epoch, we drew 5000 samples from the training data using random sampling with replacement to train our model. The training process lasted at most 15 epochs and we performed early-stopping with patience of 4 epochs based on the validation performance, which took approximately 40 minutes on an Nvidia GeForce RTX 3090 GPU. In the testing phase, it took about 5 seconds to make prediction for one protein with pre-computed features.

### 2.5 Evaluation metrics

Similar to the previous studies [56, 57], we used specificity (Spe), precision (Pre), recall (Rec), F1-score (F1), Matthews correlation coefficient (MCC), area under the receiver operating characteristic curve (AUC), and area under the precision-recall curve (AUPR) to measure the predictive performance:

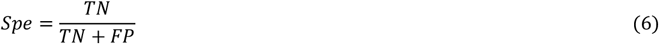

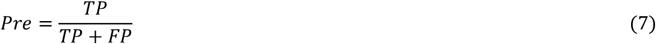

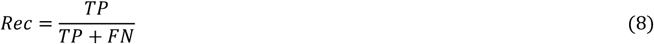

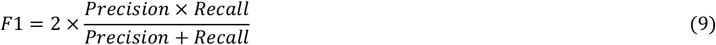

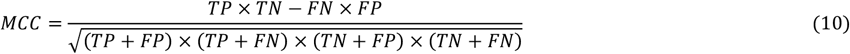

where true positives (TP) and true negatives (TN) denote the number of binding and non-binding sites identified correctly, and false positives (FP) and false negatives (FN) denote the number of incorrectly predicted binding and non-binding sites, respectively. AUC and AUPR are independent of thresholds, thus revealing the overall performance of a model. The other metrics were calculated using a threshold to convert the predicted binding probabilities to binary predictions, which was determined by maximizing F1-score for the model. We used AUPR for the above hyperparameter selection as it is more sensitive and it emphasizes more on the minority class in imbalanced two-class classification tasks [58].

Significance tests were performed to investigate if the results are not biased by a subset of test proteins by measuring whether the predictive performance is consistent over different subsets. Similar to [26, 59], we randomly sampled 70% of the test proteins and calculated the AUPRs of the best-performing method and other methods. This was repeated 10 times and we compared the corresponding 10 paired results. If the measurements were normal, as tested by the Anderson–Darling test [60] with 0.05 significance, we applied the paired t-test to investigate significance. Otherwise, the Wilcoxon rank sum test [61] was utilized. If *P*-value < 0.05, the difference between a given pair of methods is considered statistically significant.

## 3. Results

### 3.1 Performance on the 5-fold cross-validation and independent tests

We evaluated the performance of GraphSite by AUC and AUPR using 5-fold cross-validation (CV) on the Train_573 dataset and independent tests on the Test_129 and Test_181 datasets. The final GraphSite model obtains AUC of 0.915, 0.934 and 0.917; as well as AUPR of 0.589, 0.544 and 0.369 on the 5-fold CV and two independent tests, respectively. The consistent AUC on the CV and tests indicate the robustness of our model. Note that the AUPR drops about 0.18 in Test_181, which might be ascribed to the lower positive sample ratio, or the lower predicted qualities of the protein structures in this dataset (discussed in **Section 3.4**). Since the prediction of GraphSite is the average predictive scores from the five trained models in cross-validation, we also discussed the uncertainty of our method, which is empirically measured by the standard deviation of these five scores. As shown in **Supplementary Table S2**, the predictions are more accurate when GraphSite is more confident and vice versa.

In order to demonstrate the advantages of protein geometric knowledge and the graph transformer model, we compared GraphSite with a baseline method BiLSTM, which contains a two-layer bidirectional long short-term memory network with 256 hidden units and an MLP module. This model uses the same residue features as GraphSite and serves as a geometric-agnostic baseline to evaluate the impact of the spatial information for binding residue prediction. As shown in **Supplementary Table S3**, GraphSite yields higher F1, MCC, AUC, and AUPR values, which are 0.064(0.051), 0.068(0.054), 0.021(0.036) and 0.073(0.073) higher than those of BiLSTM on Test_129(Test_181), respectively. **Figure 2** and **Supplementary Figure S1** show the precision-recall curves and ROC curves of GraphSite and BiLSTM on the CV, Test_129 and Test_181, where the curves of GraphSite are largely located above those of BiLSTM. Here, the *k*-nearest mask helps GraphSite focus on the spatially adjacent residues, while remote residues can still be learned since the whole graph is connected. As shown in **Supplementary Table S4**, removal of the *k*-nearest mask leads to AUPR drops (0.036 and 0.034) on both Test_129 and Test_181.

**Figure 2.**
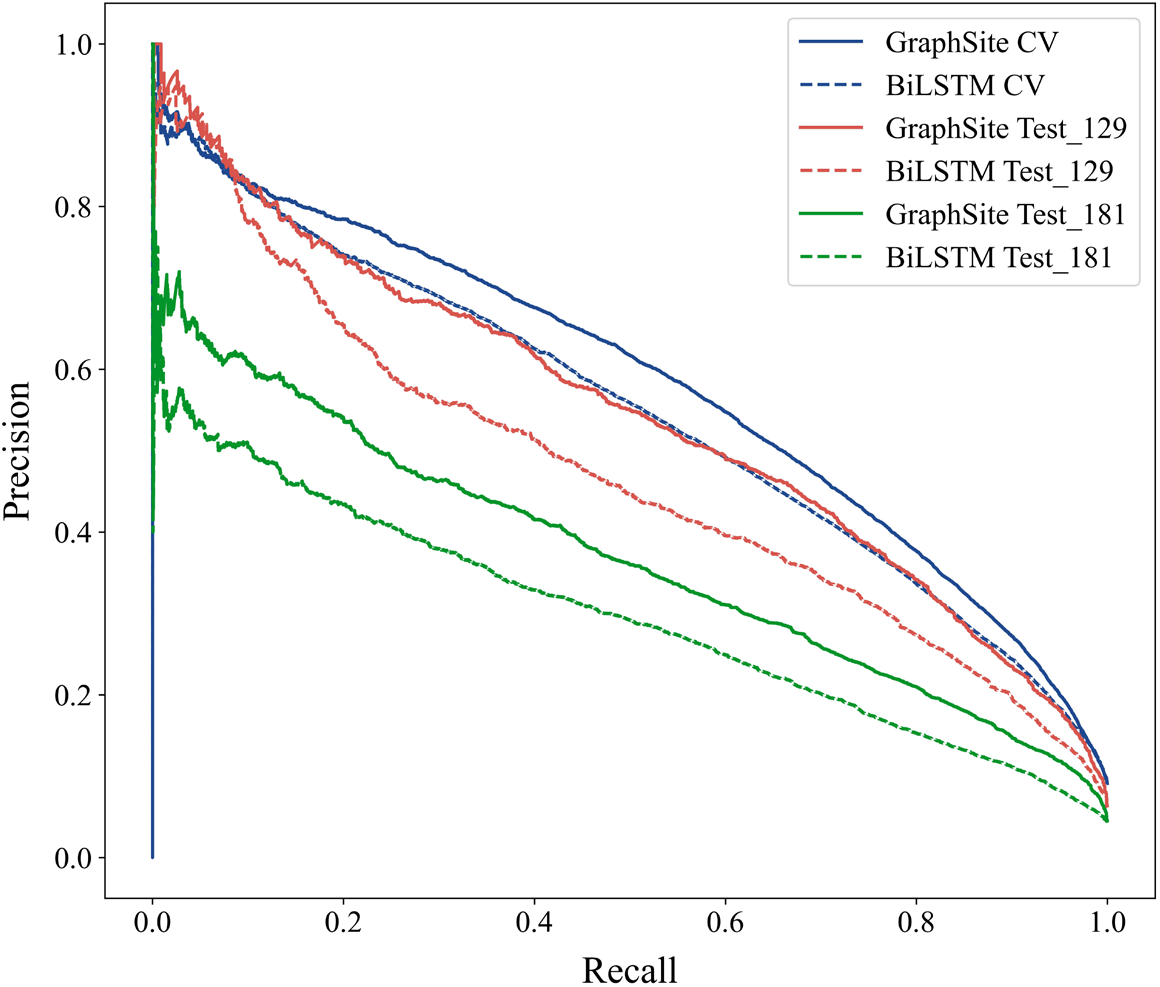
Precision-recall curves of GraphSite and BiLSTM on the CV, Test_129 and Test_181.

The performance improvement of GraphSite over BiLSTM is likely due to its better capability in capturing long-range contact information. To illustrate this, we compared the performance of GraphSite and BiLSTM on amino acids with different number of non-local contacts, defined as the contacts from the residues that are more than 20 residues away in sequence positions, but ≤ 12 Å in terms of their atomic distances between Cα atoms. **Figure 3** shows that GraphSite consistently surpasses BiLSTM on Test_181 and more importantly, the performance gap between them enlarges as the non-local contact number of the amino acids increases. Specifically, the performance of GraphSite surpasses BiLSTM by 8.6% in MCC on the amino acids with 0 to 9 non-local contacts, and the gap widens to as much as 80.5% on the amino acids with ≥ 30 non-local contacts. Similar results can also be observed in Test_129 (**Supplementary Figure S2**). These comparisons highlight the importance of the spatial information, and the effectiveness of GraphSite in harnessing protein structural knowledge especially non-local contacts for DNA-binding residue recognition.

**Figure 3.**
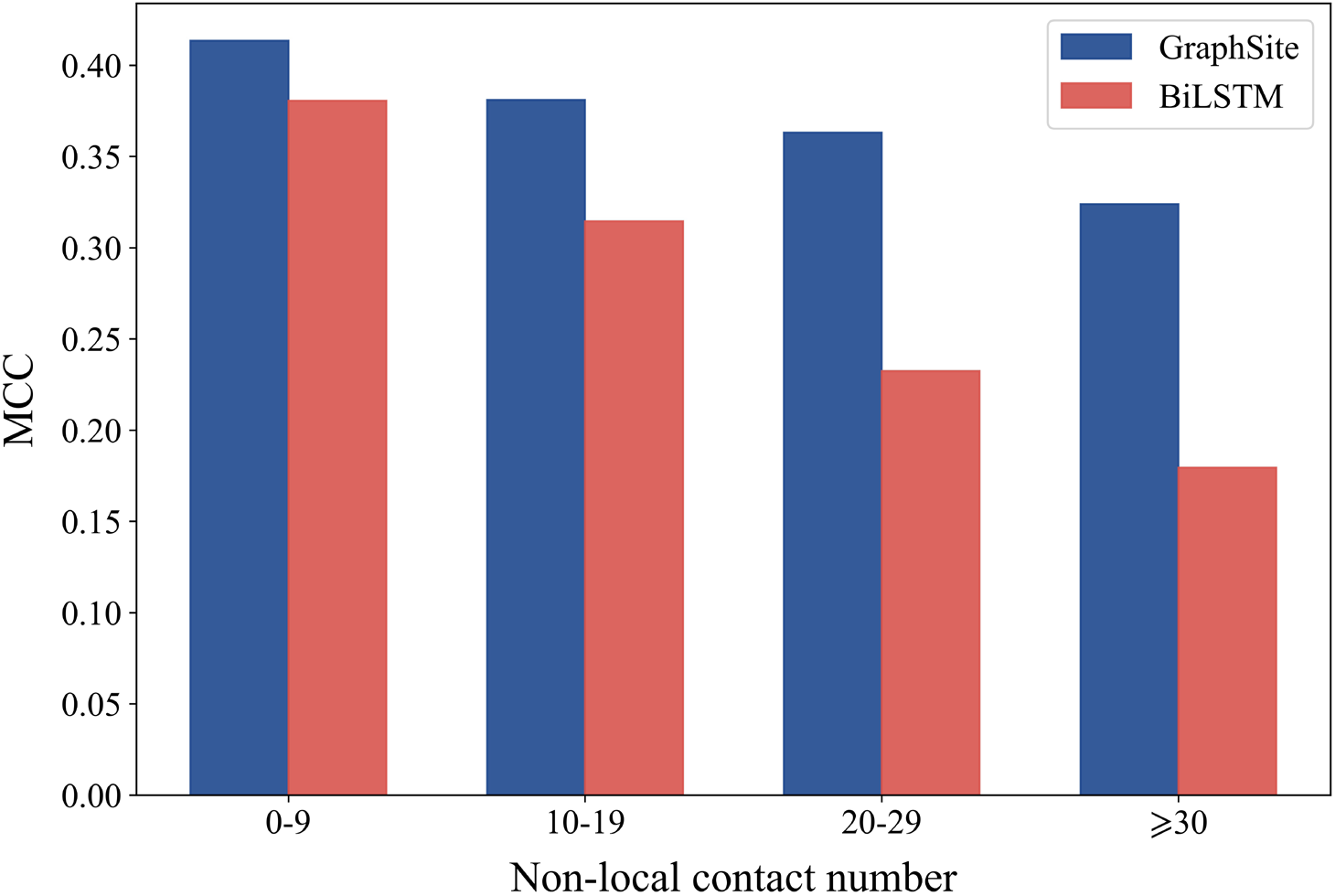
The MCC of GraphSite and BiLSTM on amino acids with different number of non-local contacts in Test_181.

### 3.2 Feature importance

To demonstrate the relative importance of each feature we adopted, we conducted feature ablation experiments by using one feature individually or excluding one feature from the final feature combination. As shown in **Table 2**, when using single feature group as node features, the highly processed single representation from AlphaFold2 already gives satisfactory performance with AUPR of 0.520 in Test_129 and 0.336 in Test_181. On the other hand, using DSSP solely as node features yields the worst performance with AUPR of 0.126 in Test_129 and 0.080 in Test_181, indicating that structural properties of amino acids such as secondary structure and relative solvent accessibility are insufficient to capture the complicated patterns of DNA-binding sites. Vice versa, the removal of single representation from the final feature combination leads to the greatest performance drops of about 0.1 in AUPR for the two test sets, and the removal of DSSP leads to the smallest performance drops as expected. In addition, although both the single representation and evolutionary features (PSSM and HMM) contain MSA information, the removal of PSSM and HMM still leads to AUPR drop of 0.021 in Test_129 and 0.037 in Test_181. The performance reduction when removing any feature group suggests that the combined feature groups are nonredundant.

**Table 2.** The AUC and AUPR of the 5-fold cross-validation (CV), Test_129 and Test_181 using a single feature individually or excluding each feature in turn from the final feature combination. AF2 Single denotes the single representation produced by AlphaFold2, and Evo denotes the evolutionary feature groups PSSM and HMM. Bold fonts indicate the best results.

| Feature | CV AUC | CV AUPR | Test_129 AUC | Test_129 AUPR | Test_181 AUC | Test_181 AUPR |
| --- | --- | --- | --- | --- | --- | --- |
| AF2 Single | 0.906 | 0.557 | 0.925 | 0.520 | 0.908 | 0.336 |
| Evo | 0.864 | 0.417 | 0.888 | 0.381 | 0.854 | 0.232 |
| DSSP | 0.691 | 0.173 | 0.721 | 0.126 | 0.686 | 0.080 |
| -AF2 Single | 0.886 | 0.497 | 0.913 | 0.452 | 0.882 | 0.263 |
| -Evo | 0.911 | 0.568 | 0.928 | 0.523 | 0.909 | 0.332 |
| -DSSP | 0.911 | 0.573 | 0.929 | 0.532 | 0.909 | 0.347 |
| GraphSite | **0.915** | **0.589** | **0.934** | **0.544** | **0.917** | **0.369** |

### 3.3 Comparison with state-of-the-art methods

We compared GraphSite with three sequence-based (SVMnuc, NCBRPred and DNAPred) and four structure-based (COACH-D, NucBind, DNABind and GraphBind) predictors on Test_129, where GraphSite outperforms all other methods significantly (shown in **Table 3**). Concretely, GraphSite surpasses the second-best sequence-based method DNAPred by 56.3% in MCC and 48.2% in AUPR, respectively. In addition, GraphSite achieves recall and precision of 0.665 and 0.460 on Test_129, respectively, surpassing all other sequence-based methods. It should be noted that recall and precision are unbalanced measures strongly depending on thresholds. Though GraphSite is a sequence-based predictor input with protein sequences only, GraphSite outperforms the latest structure-based method GraphBind by 0.020 in MCC and 0.025 in AUPR, respectively. This is reasonable because: 1) GraphBind only uses the evolutionary features PSSM and HMM from MSA, while our method additionally employs the informative single representation from AlphaFold2. 2) The graph transformer model is proven to be powerful (**Figure 2**). 3) The AlphaFold2-predicted protein structures used by GraphSite are of high quality (discussed in **Section 3.4**). On the other hand, the performance of these four structure-based methods will further decrease when using predicted structures as input (e.g., AUPR from 0.519 to 0.497 for GraphBind), and the superiority of our method will be further reflected. Besides, we also tested GraphSite using the native structures as input, which will cause performance drop of AUPR from 0.544 to 0.502, since our method were trained using predicted structures. However, re-training the model using native structures will restore the AUPR to a similar level (0.541) as before.

**Table 3.** Performance comparison of GraphSite with state-of-the-art methods on Test_129 and independent Test_181. The results of GraphBind were obtained from its standalone program, while the predictions by other competitive methods were generated from their web servers. Bold fonts indicate the best results.

| Dataset | Method | Spe | Rec | Pre | F1 | MCC | AUC | AUPR | *P*-values of AUPR |
| --- | --- | --- | --- | --- | --- | --- | --- | --- | --- |
| Test_129 | SVMnuc | 0.966 | 0.316 | 0.371 | 0.341 | 0.304 | 0.812 | 0.302 | $6.6 \times 10^{-14}$ |
| | NCBRPred | 0.969 | 0.312 | 0.392 | 0.347 | 0.313 | 0.823 | 0.310 | $2.2 \times 10^{-11}$ |
| | DNAPred | 0.954 | 0.396 | 0.353 | 0.373 | 0.332 | 0.845 | 0.367 | $2.3 \times 10^{-11}$ |
| | COACH-D<sup>a</sup> | 0.958 | 0.367 | 0.357 | 0.362 | 0.321 | 0.710 | 0.269 | $2.7 \times 10^{-11}$ |
| | NucBind<sup>a</sup> | 0.966 | 0.330 | 0.381 | 0.354 | 0.317 | 0.811 | 0.294 | $9.2 \times 10^{-12}$ |
| | DNABind<sup>a</sup> | 0.926 | 0.601 | 0.346 | 0.440 | 0.411 | 0.858 | 0.402 | $1.9 \times 10^{-10}$ |
| | GraphBind<sup>a</sup> | 0.941 | 0.684 | 0.422 | 0.522 | 0.500 | 0.928 | 0.519 | $6.1 \times 10^{-5}$ |
| | COACH-D<sup>b</sup> | 0.955 | 0.328 | 0.318 | 0.323 | 0.279 | 0.712 | 0.248 | $3.1 \times 10^{-11}$ |
| | NucBind<sup>b</sup> | 0.964 | 0.322 | 0.366 | 0.343 | 0.304 | 0.809 | 0.284 | $8.0 \times 10^{-13}$ |
| | DNABind<sup>b</sup> | 0.952 | 0.487 | 0.389 | 0.433 | 0.395 | 0.832 | 0.391 | $1.1 \times 10^{-11}$ |
| | GraphBind<sup>b</sup> | 0.948 | 0.625 | 0.434 | 0.512 | 0.484 | 0.916 | 0.497 | $2.6 \times 10^{-7}$ |
|  | GraphSite | 0.950 | 0.665 | 0.460 | **0.543** | **0.519** | **0.934** | **0.544** | N/A |
| Test_181 | NCBRPred | 0.964 | 0.259 | 0.241 | 0.250 | 0.215 | 0.771 | 0.183 | $7.2 \times 10^{-11}$ |
| | SVMnuc | 0.960 | 0.289 | 0.242 | 0.263 | 0.229 | 0.803 | 0.193 | $7.1 \times 10^{-11}$ |
| | DNAPred | 0.948 | 0.334 | 0.223 | 0.267 | 0.233 | 0.802 | 0.230 | $6.2 \times 10^{-10}$ |
| | COACH-D<sup>a</sup> | 0.971 | 0.254 | 0.280 | 0.266 | 0.235 | 0.655 | 0.172 | $2.8 \times 10^{-12}$ |
| | NucBind<sup>a</sup> | 0.960 | 0.293 | 0.248 | 0.269 | 0.234 | 0.796 | 0.191 | $7.6 \times 10^{-11}$ |
| | DNABind<sup>a</sup> | 0.904 | 0.535 | 0.199 | 0.290 | 0.279 | 0.825 | 0.219 | $4.6 \times 10^{-10}$ |
| | GraphBind<sup>a</sup> | 0.933 | 0.624 | 0.293 | 0.399 | 0.392 | 0.904 | 0.339 | $3.9 \times 10^{-5}$ |
| | COACH-D<sup>b</sup> | 0.971 | 0.239 | 0.266 | 0.251 | 0.220 | 0.668 | 0.169 | $3.4 \times 10^{-12}$ |
| | NucBind<sup>b</sup> | 0.959 | 0.288 | 0.240 | 0.262 | 0.227 | 0.798 | 0.186 | $5.5 \times 10^{-11}$ |
| | DNABind<sup>b</sup> | 0.941 | 0.392 | 0.229 | 0.289 | 0.259 | 0.803 | 0.208 | $2.8 \times 10^{-10}$ |
| | GraphBind<sup>b</sup> | 0.949 | 0.505 | 0.304 | 0.380 | 0.357 | 0.893 | 0.317 | $7.4 \times 10^{-7}$ |
|  | GraphSite | 0.958 | 0.517 | 0.354 | **0.420** | **0.397** | **0.917** | **0.369** | N/A |

<sup>a</sup> Using native protein structures.
<sup>b</sup> Using predicted protein structures.

To further demonstrate the generalization and stability of our method, we also compared GraphSite with other methods on our newly built independent Test_181. Note that this is a more challenging dataset for our method since we set “--max_template_date” in AlphaFold2 before the release dates of all proteins in Test_181. As shown in **Table 3**, the performance ranks of these methods are generally consistent as in Test_129, and GraphSite still outperforms all other methods significantly, including the structure-based methods that use native protein structures. On the other hand, when using predicted structures as input, our method surpasses the best structure-based method GraphBind by 16.4% in AUPR and 11.2% in MCC, respectively. This suggests that our method is practical and much more powerful for the situation where only protein sequences instead of native structures are available.

### 3.4 Impact of the quality of predicted protein structure

Since GraphSite employs predicted protein structures for geometric deep learning, the predicted quality of AlphaFold2 should be crucial to the downstream DNA-binding site prediction. We calculated the average global distance test (GDT) [62] between the native structures and the predicted structures by AlphaFold2 in the three datasets through SPalign [63]. The average GDT for Train_573 and Test_129 are 0.86 and 0.85, indicating that AlphaFold2 can make accurate structure predictions when similar structure templates are available. However, the predicted quality drops when we set the restriction of the max available date of templates in AlphaFold2. Concretely, the GDT drops to 0.71 for the independent Test_181, which partly explains why the performance of GraphSite decreases in this dataset. For further demonstration, **Figure 4** shows the predicted quality (measured by GDT) of AlphaFold2 and the per-protein AUPR on Test_181 for GraphSite (blue scatters). Moreover, we sorted the proteins in Test_181 according to the GDT and divided them into 9 bins equally to compute the average GDT and AUPR for each bin (red line). As expected, GraphSite shows a positive correlation between the predicted quality of AlphaFold2 and the AUPR. A closer inspection shows that the top-30% proteins with the highest GDT (average GDT = 0.92) correspond to an average AUPR of 0.525 predicted by GraphSite. On the other hand, the bottom-30% proteins with the lowest GDT (average GDT = 0.46) correspond to an average AUPR of 0.287, which is significantly lower than that of the top-30% proteins (*P*-value = 9.0 × 10^−5^) according to Mann–Whitney *U* test [64]. We also observed a negative correlation between the predicted error of AlphaFold2 at amino acid level (measured by the distance between the native and predicted amino acid after structure alignment) and the performance of GraphSite (**Supplementary Table S5**). These results suggest the importance of the predicted quality of protein structure for DNA-binding site prediction.

**Figure 4.**
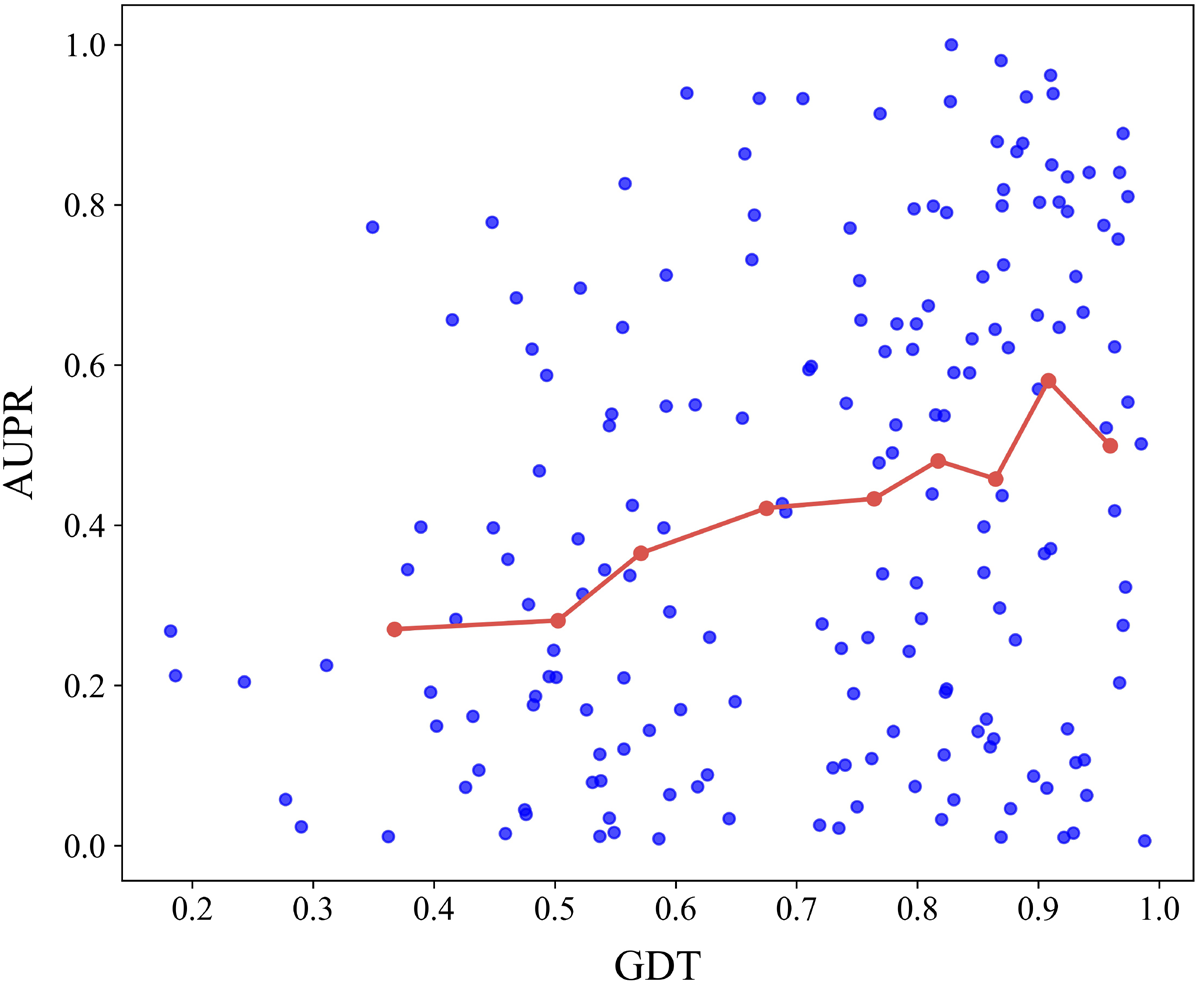
The positive correlation between the predicted quality of AlphaFold2 measured by GDT and the performance of GraphSite measured by AUPR on Test_181. The blue scatters denote the per-protein GDT and AUPR, while the red line denotes the average GDT and AUPR for each bin after sorting all proteins according to GDT and dividing them into 9 bins.

### 3.5 GraphSite learns effective latent representations of residues

In this section, we visualized the raw feature vectors and the latent feature vectors learned by GraphSite on Test_181. For a target residue, the initial node feature vector consisting of MSA information and structural properties with the size of 438 serves as the raw feature vector. The latent feature vector learned by GraphSite with the size of 128 is the concatenation of the embedding vectors from the two graph transformer layers. t-SNE [65] was applied to project the high-dimensional feature vectors into the 2-dimensional space. **Figure 5A** and **5B** illustrate the distributions of samples encoded by raw feature vectors and latent feature vectors, respectively. As shown in **Figure 5A**, the binding and non-binding residues overlap and are indistinguishable, while **Figure 5B** shows that most binding residues are clustered together and separated from most non-binding residues. These results demonstrate that the latent representations learned by GraphSite effectively improve the discriminability of binding and non-binding residues.

**Figure 5.**
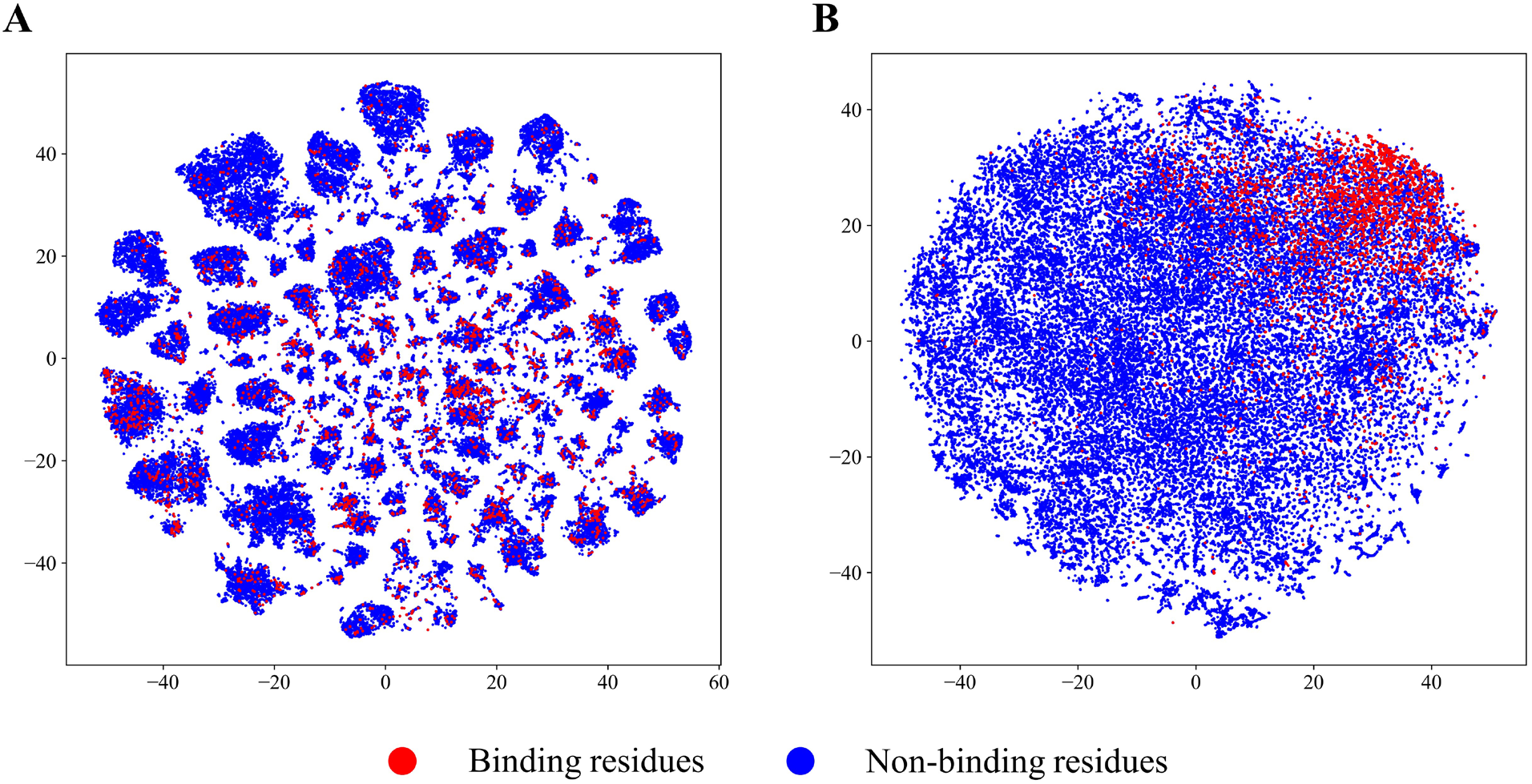
Visualization of the distributions of samples encoded by raw feature vectors (**A**) and latent feature vectors learned by GraphSite (**B**) on Test_181 using t-SNE.

### 3.6 Case study

To visualize the superiority of our method, we selected mycobacterial DNA polymerase LigD (PDB ID: 6SA0, chain A) from Test_181 for illustration. **Figure 6** shows the DNA-binding site prediction results of GraphSite (**A**) and the geometric-agnostic baseline method BiLSTM (**B**). In this example, there are 32 DNA-binding residues over a total of 333 residues. GraphSite predicts 48 binding residues in which 24 are true positives, leading to an F1 of 0.600, MCC of 0.562, and AUPR of 0.554. By comparison, BiLSTM predicts 52 binding residues in which only 20 are true positives, leading to a lower F1 of 0.476, MCC of 0.421, and AUPR of 0.367. Besides, as shown in **Figure 6A**, the false-positive binding residues (colored in red) predicted by GraphSite are mostly around the interface of protein-DNA interaction or close to the DNA structure. In addition, the results of the structure-based method GraphBind using the predicted structure of this protein can be found in **Supplementary Figure S3.** Visualization of another case (PDB ID: 6YMW, chain B) can also be found in **Supplementary Figure S4**.

**Figure 6.**
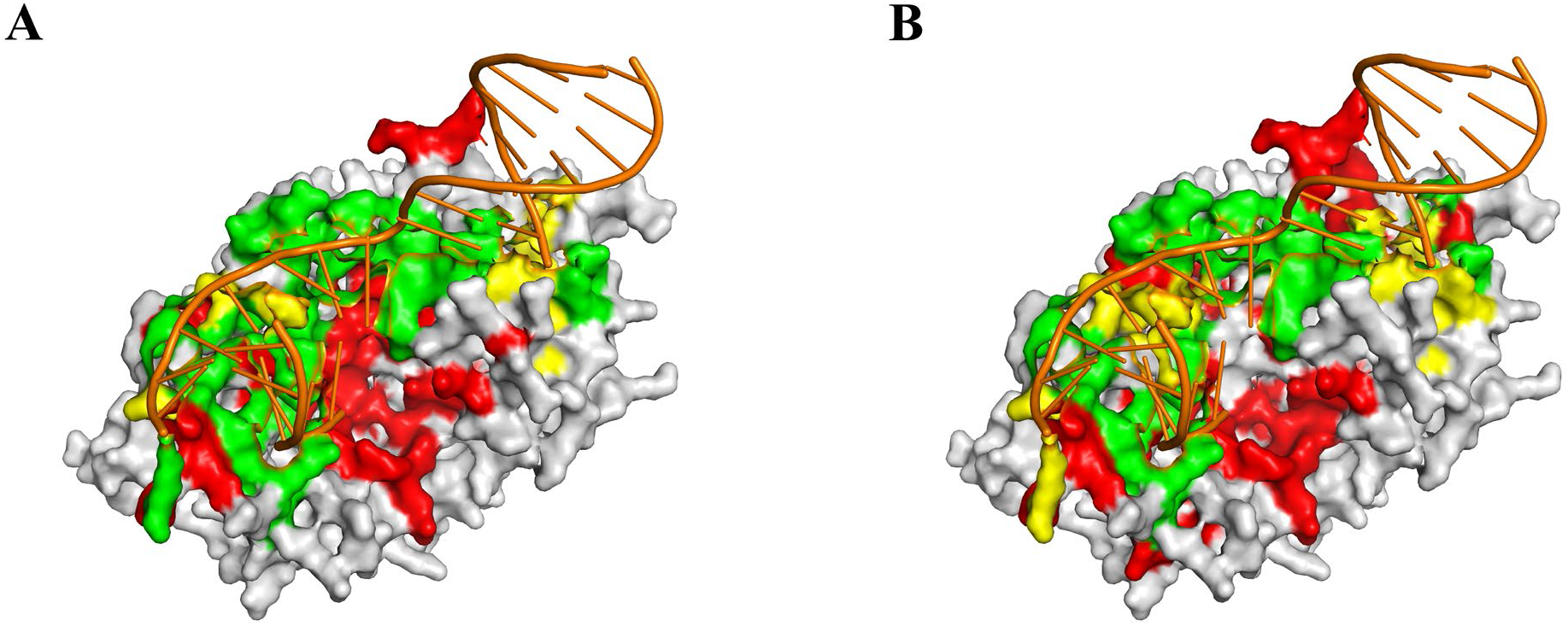
Visualization of one example (PDB ID: 6SA0, chain A) from Test_181 predicted by GraphSite (**A**) and the geometric-agnostic baseline method BiLSTM (**B**). True positives, false positives and false negatives are colored in green, red and yellow, respectively.

## 4. Discussion and Conclusion

Identifying protein-DNA binding sites is crucial for understanding biological activities and designing novel drugs. Existing sequence-based methods only consider contextual features of the sequential neighbors, leading to their limited predictive performance, while the structure-based methods are not applicable to most proteins that don’t have known tertiary structures. Trained with the predicted structure models and the single representations from AlphaFold2, GraphSite achieves great performance (surpassing the best structure-based method) using only protein sequences, which simultaneously solves the limitations of the current sequence-based and structure-based methods. The graph transformer technique adopted by GraphSite is able to refine the geometric characteristics by taking the local structural context topology into account, while most of the competitive methods first extract structural characteristics and then feed these features into some supervised classifiers, separating the feature engineering and classification. In summary, the superiority of GraphSite benefits from two aspects: 1) the predicted structures from AlphaFold2 are of high quality and the single representations are informative; 2) the structure-aware graph transformer is an effective algorithm to learn the patterns for binding residue prediction.

With the development of sequencing techniques, many DNA-binding proteins have been detected, and such discoveries require confirmations in biological experiments. While the whole chain screening is time-consuming and expensive, the predictive methods can help narrow down potential binding sites, as indicated in our previous collaborative study [10] to validate nucleic acid binding residues in JAK2 kinase through computational predictions and wet experiments. On the other hand, these predictions can also provide hypotheses and insights for the mechanisms of many disease-causing gene mutations, such as the THOC2 mutations which affect mRNA export [9]. In novel drug design, the binding site prediction can be used to predict druggability [13] or used as condition of generative models for de novo molecule design [14].

Our GraphSite model still has a few limitations. Firstly, the performance of GraphSite is largely affected by the predicted quality of AlphaFold2. This may be solved by adding other informative sequence-derived features or building heterogeneous graphs through integrating protein primary sequences to increase the model robustness to structure predicted quality. Another way is to keep all five relaxed models from AlphaFold2 instead of only retaining the model with the highest predicted LDDT score to perform data augmentation. Secondly, our graph transformer algorithm only uses the protein distance matrices as masks in the self-attention step now, and modifications might be made to add the pairwise residue distances as biases to the attention scores. Our framework can also be explored to handle protein graphs with edge features constructed by residue distance, angle or the pair representation from AlphaFold2. Thirdly, our method only considers the protein information to predict potential DNA-binding sites, thus cannot predict the specific binding pattern given a known DNA sequence or structure. We left these above improvements of this longstanding challenge to future work.

In conclusion, this study proposes a geometric-aware framework called GraphSite for DNA-binding site prediction, where we predict protein structures from sequences using AlphaFold2 and employ graph transformer network to learn the amino acid representations. GraphSite shows preferable performance than other sequence-based and structure-based methods in comprehensive evaluations. We suggest that our method could provide useful information for biologists studying protein-DNA binding patterns or pathogenic mechanisms of mutations, and chemists interested in targeted drug design. In the future, we would further improve our graph transformer architecture and integrate multi-task learning [66] to extend our method to various fields, including predicting protein binding sites with RNA and small ligands, or protein functional sites such as methylation and phosphorylation.

## Key points

- Existing sequence-based methods for identifying protein-DNA binding sites only consider contextual features of the sequential neighbors, which are limited to capture spatial information.
- GraphSite is the first sequence-based method to predict protein-DNA binding sites based on the predicted structures from AlphaFold2, where structure-aware graph transformer is employed to capture the protein structural context topology.
- GraphSite shows preferable performance than state-of-the-art sequence-based and structure-based methods in two independent datasets.

## Data Availability

We provide the datasets, the predicted structures, and the source codes along with the pre-trained models of GraphSite at https://github.com/biomed-AI/GraphSite. The GraphSite web server is freely available at https://biomed.nscc-gz.cn/apps/GraphSite.

## Supplementary data

Supplementary data are available online at https://academic.oup.com/bib. We also provide the datasets, the predicted structures, and the source codes along with the pre-trained models of GraphSite at https://github.com/biomed-AI/GraphSite.

## Acknowledgments

The authors are very grateful to Wei Lu and Jixian Zhang from Galixir Technologies Ltd for contributing to the training code of GraphSite.

## Supplementary Information

**Table S1.** The performance of GraphSite on the 5-fold cross-validation (CV), Test_129 and Test_181 using different hyperparameters. For each row, the hyperparameters that are not shown are the same as the final GraphSite model.

| Hyperparameter | Value | 5-Fold CV AUPRC | Test_129 AUPRC | Test_181 AUPRC |
| --- | --- | --- | --- | --- |
| Layers | 1 | 0.570 | 0.524 | 0.356 |
|  | **2** | 0.589 | 0.544 | 0.369 |
|  | 4 | 0.577 | 0.530 | 0.359 |
| *k* | 20 | 0.573 | 0.518 | 0.356 |
|  | **30** | 0.589 | 0.544 | 0.369 |
|  | 40 | 0.569 | 0.522 | 0.352 |
| *h* | 1 | 0.548 | 0.506 | 0.337 |
|  | 2 | 0.578 | 0.532 | 0.362 |
|  | **4** | 0.589 | 0.544 | 0.369 |
|  | 8 | 0.583 | 0.536 | 0.353 |
| Hidden Units | 32 | 0.580 | 0.525 | 0.359 |
|  | **64** | 0.589 | 0.544 | 0.369 |
|  | 128 | 0.549 | 0.500 | 0.333 |
| Dropout | 0.1 | 0.569 | 0.520 | 0.352 |
|  | **0.2** | 0.589 | 0.544 | 0.369 |
|  | 0.3 | 0.588 | 0.530 | 0.363 |

**Table S2.** The negative correlation between the predictive uncertainty and the performance of GraphSite on Test_129 and Test_181. Amino acids in the datasets were sorted according to the uncertainty (measured by the standard deviation of the five predictive scores), and then divided into 5 bins equally. Average uncertainty, accuracy (ACC) and binary cross-entropy (BCE) were calculated for each bin.

| Dataset | Bin | 1 | 2 | 3 | 4 | 5 |
| --- | --- | --- | --- | --- | --- | --- |
| Test_129 | Average Uncertainty | $4.6 \times 10^{-4}$ | $1.2 \times 10^{-3}$ | $3.7 \times 10^{-3}$ | $1.8 \times 10^{-2}$ | $1.3 \times 10^{-1}$ |
|  | ACC | 0.999 | 0.998 | 0.992 | 0.965 | 0.712 |
|  | BCE | 0.003 | 0.013 | 0.046 | 0.143 | 0.451 |
| Test_181 | Average Uncertainty | $4.8 \times 10^{-4}$ | $1.3 \times 10^{-3}$ | $3.8 \times 10^{-3}$ | $1.7 \times 10^{-2}$ | $1.2 \times 10^{-1}$ |
|  | ACC | 0.999 | 0.998 | 0.994 | 0.969 | 0.735 |
|  | BCE | 0.005 | 0.010 | 0.036 | 0.128 | 0.422 |

**Table S3.**
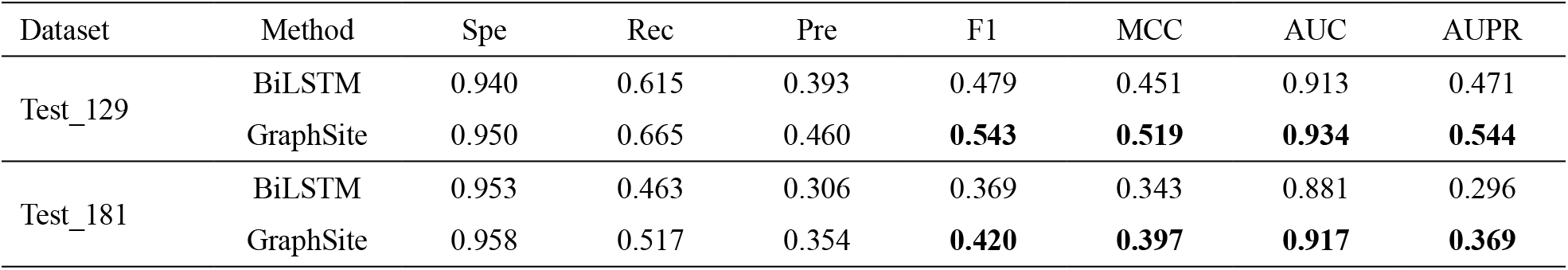
Performance comparison with the geometric-agnostic baseline model BiLSTM on Test_129 and Test_181. The highest values are bolded.

**Figure S1.**
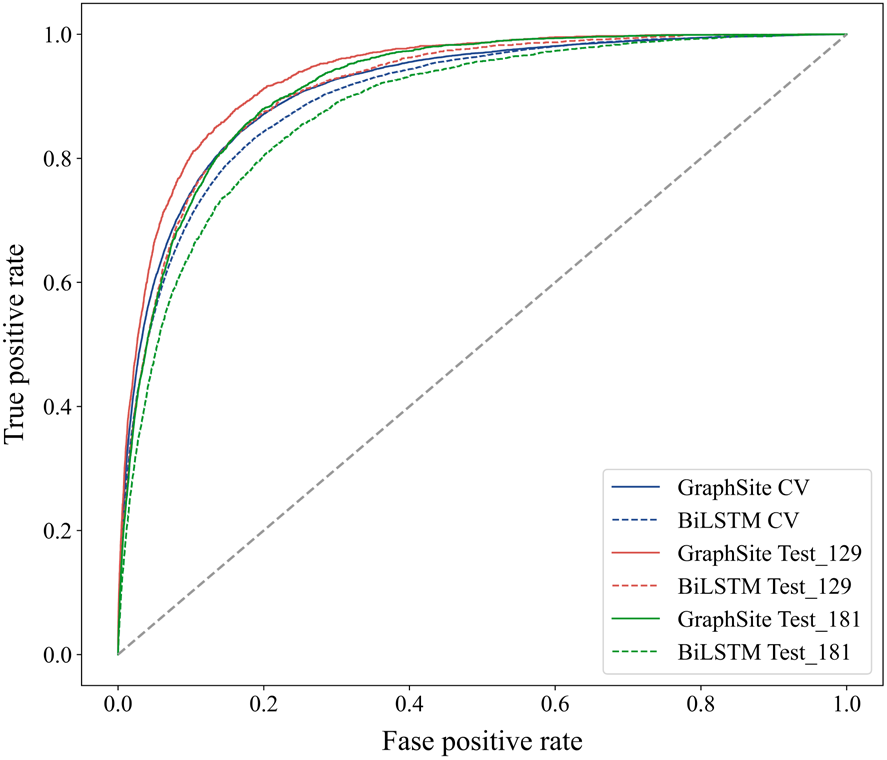
Receiver operating characteristic curves of GraphSite and BiLSTM on the CV, Test_129 and Test_181.

**Table S4.** Performance comparison with transformer without *k*-nearest mask on Test_129 and Test_181. The highest values are bolded.

| Dataset | Method | Spe | Rec | Pre | F1 | MCC | AUC | AUPR |
| --- | --- | --- | --- | --- | --- | --- | --- | --- |
| Test_129 | Transformer | 0.960 | 0.557 | 0.470 | 0.510 | 0.478 | 0.916 | 0.508 |
|  | GraphSite | 0.950 | 0.665 | 0.460 | **0.543** | **0.519** | **0.934** | **0.544** |
| Test_181 | Transformer | 0.959 | 0.464 | 0.339 | 0.392 | 0.365 | 0.896 | 0.335 |
|  | GraphSite | 0.958 | 0.517 | 0.354 | **0.420** | **0.397** | **0.917** | **0.369** |

**Figure S2.**
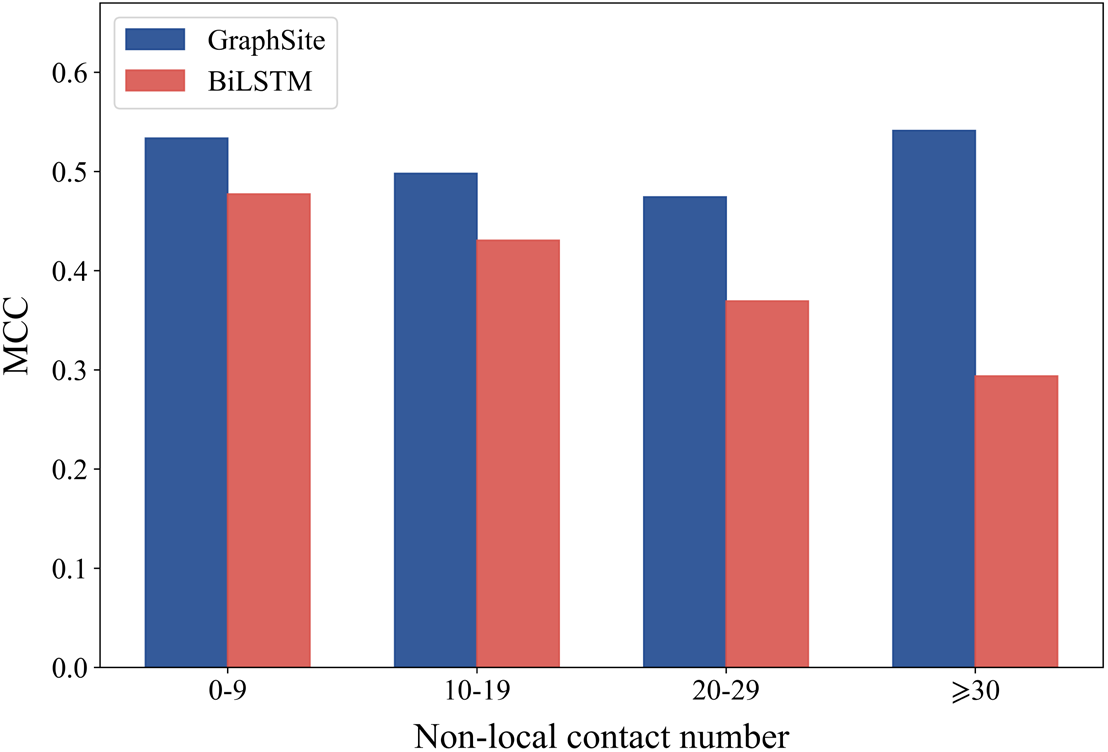
The MCC of GraphSite and BiLSTM on amino acids with different number of non-local contacts in Test_129.

**Table S5.** The model performance for the amino acids with different degrees of predicted errors. All amino acids in Test_181 were sorted according to the distance between the native and predicted amino acid and divided into 5 bins equally, from which the median distances and AUPRs were calculated.

| Bin Range (Å) | 0.00-0.60 | 0.60-1.06 | 1.06-1.93 | 1.93-6.53 | > 6.53 |
| --- | --- | --- | --- | --- | --- |
| Median Distance (Å) | 0.39 | 0.81 | 1.40 | 3.12 | 21.56 |
| AUPR | 0.445 | 0.456 | 0.347 | 0.310 | 0.258 |

**Figure S3** shows the DNA-binding site prediction results of GraphSite (**A**) and the second-best method GraphBind (**B**) using the predicted structure of mycobacterial DNA polymerase LigD (PDB ID: 6SA0, chain A) from Test_181. In this example, there are 32 DNA-binding residues over a total of 333 residues. GraphSite predicts 48 binding residues in which 24 are true positives, leading to an F1 of 0.600, MCC of 0.562, and AUPR of 0.554. By comparison, GraphBind predicts 15 binding residues in which 9 are true positives, and 72% of the true binding residues are incorrectly predicted as non-binding residues, leading to a lower F1 of 0.383, MCC of 0.371, and AUPR of 0.461.

**Figure S3.**
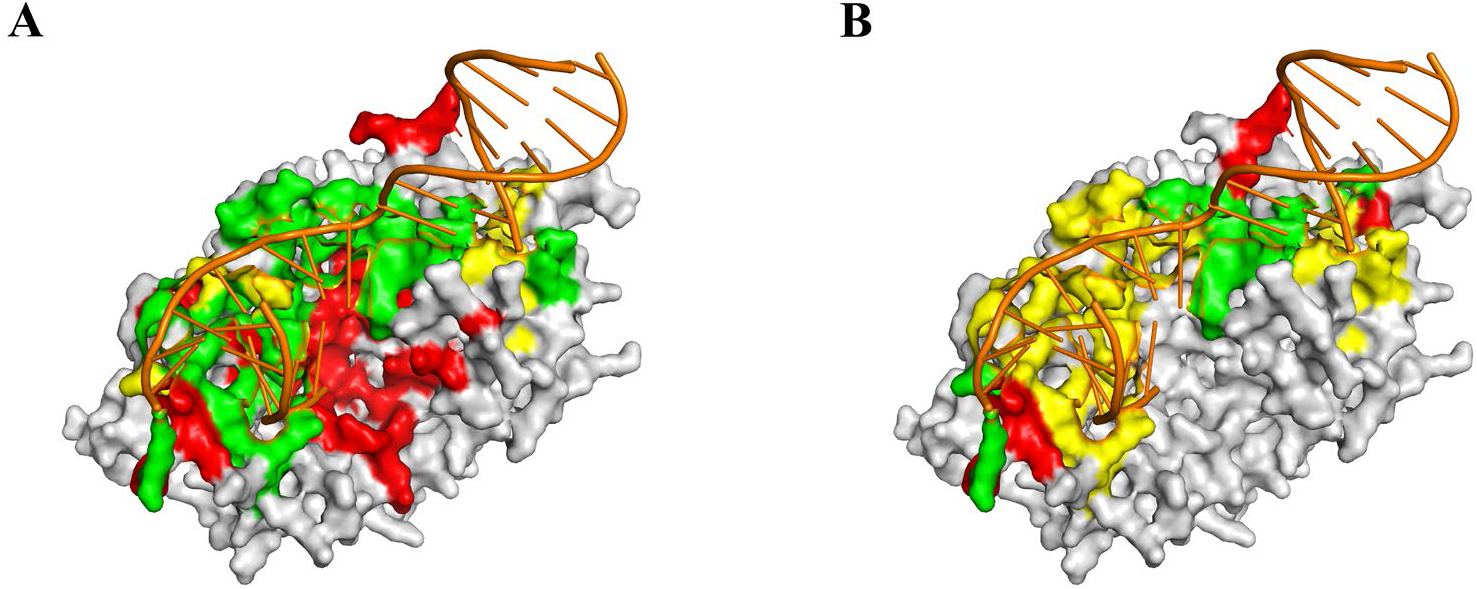
Visualization of one example (PDB ID: 6SA0, chain A) from Test_181 predicted by GraphSite (**A**) and the second-best method GraphBind (**B**). True positives, false positives and false negatives are colored in green, red and yellow, respectively.

**Figure S4** shows the DNA-binding site prediction results of GraphSite (**A**), the geometric-agnostic baseline method BiLSTM (**B**), and the second-best method GraphBind (**C**) using the predicted structure of yeast mitochondrial transcription factor 1 (PDB ID: 6YMW, chain B) from Test_181. In this example, there are 15 DNA-binding residues over a total of 340 residues. GraphSite predicts 26 binding residues in which 9 are true positives, leading to an F1 of 0.439, MCC of 0.423, and AUPR of 0.371. By comparison, BiLSTM predicts 21 binding residues in which only 4 are true positives, and 73% of the true binding residues are incorrectly predicted as non-binding residues, leading to a lower F1 of 0.222, MCC of 0.183, and AUPR of 0.273. The second-best method GraphBind predicts 15 binding residues in which 4 are true positives, leading to an F1 of 0.267, MCC of 0.233, and AUPR of 0.221.

**Figure S4.**
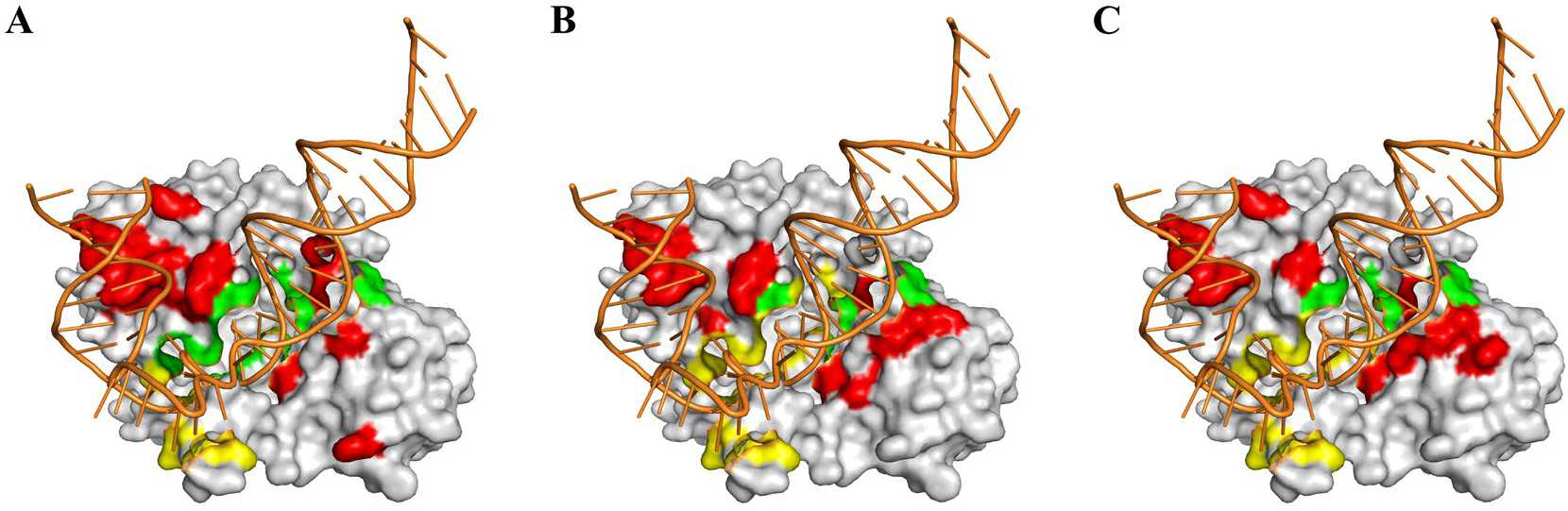
Visualization of another example (PDB ID: 6YMW, chain B) from Test_181 predicted by GraphSite (**A**), BiLSTM (**B**) and GraphBind (**C**). True positives, false positives and false negatives are colored in green, red and yellow, respectively.

## References

1. Zhao H, Yang Y, Zhou Y. Structure-based prediction of DNA-binding proteins by structural alignment and a volume-fraction corrected DFIRE-based energy function, Bioinformatics 2010;26:1857–1863.

2. Charoensawan V, Wilson D, Teichmann SA. Genomic repertoires of DNA-binding transcription factors across the tree of life, Nucleic acids research 2010;38:7364–7377.

3. Dai H, Umarov R, Kuwahara H et al. Sequence2vec: a novel embedding approach for modeling transcription factor binding affinity landscape, Bioinformatics 2017;33:3575–3583.

4. Rastogi C, Rube HT, Kribelbauer JF et al. Accurate and sensitive quantification of protein-DNA binding affinity, Proceedings of the National Academy of Sciences 2018;115:E3692–E3701.

5. Umarov R, Kuwahara H, Li Y et al. Promoter analysis and prediction in the human genome using sequence-based deep learning models, Bioinformatics 2019;35:2730–2737.

6. Yan Y, Zhang D, Zhou P et al. HDOCK: a web server for protein–protein and protein–DNA/RNA docking based on a hybrid strategy, Nucleic acids research 2017;45:W365–W373.

7. Su H, Liu M, Sun S et al. Improving the prediction of protein–nucleic acids binding residues via multiple sequence profiles and the consensus of complementary methods, Bioinformatics 2019;35:930–936.

8. Ghersi D, Sanchez R. Improving accuracy and efficiency of blind protein-ligand docking by focusing on predicted binding sites, Proteins: Structure, Function, and Bioinformatics 2009;74:417–424.

9. Kumar R, Corbett MA, Van Bon BW et al. THOC2 mutations implicate mRNA-export pathway in X-linked intellectual disability, The American Journal of Human Genetics 2015;97:302–310.

10. Wang S, Liang K, Hu Q et al. JAK2-binding long noncoding RNA promotes breast cancer brain metastasis, The Journal of clinical investigation 2017;127:4498–4515.

11. Bhardwaj N, Lu H. Residue-level prediction of DNA-binding sites and its application on DNA-binding protein predictions, FEBS letters 2007;581:1058–1066.

12. Konc J, Hodošček M, Ogrizek M et al. Structure-based function prediction of uncharacterized protein using binding sites comparison, PLoS computational biology 2013;9:e1003341.

13. Schmidtke P, Barril X. Understanding and predicting druggability. A high-throughput method for detection of drug binding sites, Journal of medicinal chemistry 2010;53:5858–5867.

14. Xu M, Ran T, Chen H. De novo molecule design through the molecular generative model conditioned by 3D information of protein binding sites, Journal of Chemical Information and Modeling 2021;61:3240–3254.

15. Orengo CA, Michie AD, Jones S et al. CATH–a hierarchic classification of protein domain structures, Structure 1997;5:1093–1109.

16. Mandel-Gutfreund Y, Margalit H. Quantitative parameters for amino acid-base interaction: implications for prediction of protein-DNA binding sites, Nucleic acids research 1998;26:2306–2312.

17. Wadkins RM. Targeting DNA secondary structures, Current medicinal chemistry 2000;7:1–15.

18. Brázda V, Hároníková L, Liao JC et al. DNA and RNA quadruplex-binding proteins, International journal of molecular sciences 2014;15:17493–17517.

19. Ahmad S, Keskin O, Sarai A et al. Protein–DNA interactions: structural, thermodynamic and clustering patterns of conserved residues in DNA-binding proteins, Nucleic acids research 2008;36:5922–5932.

20. Zhu Y-H, Hu J, Song X-N et al. DNAPred: accurate identification of DNA-binding sites from protein sequence by ensembled hyperplane-distance-based support vector machines, Journal of chemical information and modeling 2019;59:3057–3071.

21. Zhang J, Ghadermarzi S, Katuwawala A et al. DNAgenie: accurate prediction of DNA-type-specific binding residues in protein sequences, Briefings in Bioinformatics 2021;22.

22. Zhang J, Chen Q, Liu B. NCBRPred: predicting nucleic acid binding residues in proteins based on multilabel learning, Briefings in Bioinformatics 2021;22.

23. Jones S, Shanahan HP, Berman HM et al. Using electrostatic potentials to predict DNA-binding sites on DNA-binding proteins, Nucleic acids research 2003;31:7189–7198.

24. Tsuchiya Y, Kinoshita K, Nakamura H. Structure-based prediction of DNA-binding sites on proteins using the empirical preference of electrostatic potential and the shape of molecular surfaces, PROTEINS: structure, Function, and Bioinformatics 2004;55:885–894.

25. Jiménez J, Doerr S, Martínez-Rosell G et al. DeepSite: protein-binding site predictor using 3D-convolutional neural networks, Bioinformatics 2017;33:3036–3042.

26. Xia Y, Xia C-Q, Pan X et al. GraphBind: protein structural context embedded rules learned by hierarchical graph neural networks for recognizing nucleic-acid-binding residues, Nucleic acids research 2021;49:e51–e51.

27. Liu R, Hu J. DNABind: A hybrid algorithm for structure-based prediction of DNA-binding residues by combining machine learning-and template-based approaches, PROTEINS: structure, Function, and Bioinformatics 2013;81:1885–1899.

28. Wu Q, Peng Z, Zhang Y et al. COACH-D: improved protein–ligand binding sites prediction with refined ligand-binding poses through molecular docking, Nucleic acids research 2018;46:W438–W442.

29. Nagarajan R, Ahmad S, Michael Gromiha M. Novel approach for selecting the best predictor for identifying the binding sites in DNA binding proteins, Nucleic acids research 2013;41:7606–7614.

30. Jumper J, Evans R, Pritzel A et al. Highly accurate protein structure prediction with AlphaFold, Nature 2021:1–11.

31. Lam JH, Li Y, Zhu L et al. A deep learning framework to predict binding preference of RNA constituents on protein surface, Nature communications 2019;10:1–13.

32. Zheng S, Li Y, Chen S et al. Predicting drug–protein interaction using quasi-visual question answering system, Nature Machine Intelligence 2020;2:134–140.

33. Kozlovskii I, Popov P. Protein–Peptide Binding Site Detection Using 3D Convolutional Neural Networks, Journal of chemical information and modeling 2021;61:3814–3823.

34. Yuan Q, Chen J, Zhao H et al. Structure-aware protein–protein interaction site prediction using deep graph convolutional network, Bioinformatics 2021.

35. Chen J, Zheng S, Zhao H et al. Structure-aware protein solubility prediction from sequence through graph convolutional network and predicted contact map, Journal of cheminformatics 2021;13:1–10.

36. Vaswani A, Shazeer N, Parmar N et al. Attention is all you need. In: Advances in neural information processing systems. 2017, p. 5998–6008.

37. Devlin J, Chang M-W, Lee K et al. BERT: Pre-training of Deep Bidirectional Transformers for Language Understanding. In: Proceedings of the 2019 Conference of the North American Chapter of the Association for Computational Linguistics. Minneapolis, Minnesota, 2019, p. 4171–4186. Association for Computational Linguistics.

38. Zheng S, Rao J, Zhang Z et al. Predicting retrosynthetic reactions using self-corrected transformer neural networks, Journal of chemical information and modeling 2019;60:47–55.

39. Chen L, Tan X, Wang D et al. TransformerCPI: improving compound–protein interaction prediction by sequence-based deep learning with self-attention mechanism and label reversal experiments, Bioinformatics 2020;36:4406–4414.

40. Ingraham J, Garg V, Barzilay R et al. Generative Models for Graph-Based Protein Design, Advances in neural information processing systems 2019;32:15820–15831.

41. Chen J, Zheng S, Song Y et al. Learning Attributed Graph Representation with Communicative Message Passing Transformer. In: Proceedings of the Thirtieth International Joint Conference on Artificial Intelligence, {IJCAI-21}. 2021, p. 2242–2248.

42. Ying C, Cai T, Luo S et al. Do Transformers Really Perform Badly for Graph Representation? In: Thirty-Fifth Conference on Neural Information Processing Systems. Online, 2021. Curran Associates Inc., 57 Morehouse Lane, Red Hook, NY, United States.

43. Yang J, Roy A, Zhang Y. BioLiP: a semi-manually curated database for biologically relevant ligand–protein interactions, Nucleic acids research 2012;41:D1096–D1103.

44. Berman HM, Westbrook J, Feng Z et al. The protein data bank, Nucleic acids research 2000;28:235–242.

45. Fu L, Niu B, Zhu Z et al. CD-HIT: accelerated for clustering the next-generation sequencing data, Bioinformatics 2012;28:3150–3152.

46. Suzek BE, Huang H, McGarvey P et al. UniRef: comprehensive and non-redundant UniProt reference clusters, Bioinformatics 2007;23:1282–1288.

47. Mitchell AL, Almeida A, Beracochea M et al. MGnify: the microbiome analysis resource in 2020, Nucleic acids research 2020;48:D570–D578.

48. Steinegger M, Mirdita M, Söding J. Protein-level assembly increases protein sequence recovery from metagenomic samples manyfold, Nature methods 2019;16:603–606.

49. Mirdita M, von den Driesch L, Galiez C et al. Uniclust databases of clustered and deeply annotated protein sequences and alignments, Nucleic acids research 2017;45:D170–D176.

50. Steinegger M, Meier M, Mirdita M et al. HH-suite3 for fast remote homology detection and deep protein annotation, BMC bioinformatics 2019;20:1–15.

51. Altschul SF, Madden TL, Schäffer AA et al. Gapped BLAST and PSI-BLAST: a new generation of protein database search programs, Nucleic acids research 1997;25:3389–3402.

52. Remmert M, Biegert A, Hauser A et al. HHblits: lightning-fast iterative protein sequence searching by HMM-HMM alignment, Nature methods 2012;9:173–175.

53. Kabsch W, Sander C. Dictionary of protein secondary structure: pattern recognition of hydrogen-bonded and geometrical features, Biopolymers: Original Research on Biomolecules 1983;22:2577–2637.

54. Kingma DP, Ba J. Adam: A Method for Stochastic Optimization. In: 3rd International Conference on Learning Representations (Poster). 2015.

55. Paszke A, Gross S, Massa F et al. Pytorch: An imperative style, high-performance deep learning library, Advances in neural information processing systems 2019;32:8026–8037.

56. Do DT, Le TQT, Le NQK. Using deep neural networks and biological subwords to detect protein S-sulfenylation sites, Briefings in Bioinformatics 2020;22.

57. Le NQK, Ho Q-T, Nguyen T-T-D et al. A transformer architecture based on BERT and 2D convolutional neural network to identify DNA enhancers from sequence information, Briefings in Bioinformatics 2021;22.

58. Saito T, Rehmsmeier M. The precision-recall plot is more informative than the ROC plot when evaluating binary classifiers on imbalanced datasets, PloS one 2015;10:e0118432.

59. Yan J, Kurgan L. DRNApred, fast sequence-based method that accurately predicts and discriminates DNA-and RNA-binding residues, Nucleic acids research 2017;45:e84–e84.

60. Anderson TW, Darling DA. Asymptotic theory of certain” goodness of fit” criteria based on stochastic processes, The annals of mathematical statistics 1952;23:193–212.

61. Wilcoxon F. Individual Comparisons by Ranking Methods, Biometrics 1945;1:80–83.

62. Zemla A. LGA: a method for finding 3D similarities in protein structures, Nucleic acids research 2003;31:3370–3374.

63. Yang Y, Zhan J, Zhao H et al. A new size-independent score for pairwise protein structure alignment and its application to structure classification and nucleic-acid binding prediction, PROTEINS: structure, Function, and Bioinformatics 2012;80:2080–2088.

64. Mann HB, Whitney DR. On a test of whether one of two random variables is stochastically larger than the other, The annals of mathematical statistics 1947:50–60.

65. Van der Maaten L, Hinton G. Visualizing data using t-SNE, Journal of machine learning research 2008;9:2579–2605.

66. Sun Z, Zheng S, Zhao H et al. To improve the predictions of binding residues with DNA, RNA, carbohydrate, and peptide via multi-task deep neural networks, IEEE/ACM transactions on computational biology and bioinformatics 2021.

